# Developing small Cas9 hybrids using molecular modeling

**DOI:** 10.1101/2023.10.24.563270

**Authors:** Antoine Mangin, Vincent Dion, Georgina Menzies

## Abstract

The RNA-guided CRISPR-Cas9 from *Streptococcus pyogenes* is the best characterized enzyme for gene editing. Its large size, however, precludes it from being packaged together with its single guide (sg)RNA into a single adeno-associated virus, limiting *in vivo* applications. Here, we developed smaller Cas9 hybrids, made of the PAM interacting domain (PID) of *S. pyogenes* and the catalytic domains of the smaller Cas9 orthologues, as well as sgRNA cognate hybrids. Molecular modeling revealed that the presence of a sgRNA stabilizes Cas9. Making the D10A mutation to turn Cas9 into a nickase dramatically alters its binding energy to the sgRNA, showing that the approach can identify functionally relevant changes. However, we found that the four Cas9/sgRNA hybrid pairs tested in human cells failed to edit target sequences. We conclude that *in silico* approaches can identify functional changes caused by point mutations but are not sufficient for designing Cas9/sgRNA hybrids.

## Introduction

The clustered regularly interspaced short palindromic repeats (CRISPR) Cas system evolved as an immunity mechanism against phage invasion^1,2^. But once Cas9 from *Streptococcus pyogenes* was found to be a programmable RNA-guided nuclease creating DNA double-strand breaks, gene editing at high efficiency in mammalian cells became possible^3,4^. Type II CRISPR/Cas complexes are formed of a Cas9 nuclease guided to a target sequence using a CRISPR (cr)RNA. This contains a customizable sequence (∼20 nucleotides), and a trans-activating crRNA (tracrRNA) formed of one or more stem loops and are unique to each Cas9 orthologue^5^. The two RNAs can be fused together to produce a convenient single guide (sg)RNA^3^. The sgRNA-Cas9 complex recognizes its complementary sequence in genomic DNA using base pairing when the target sequence is followed by a short (3 to 8 bp), species-specific, protospacer adjacent motif (PAM). Upon binding, the HNH and RuvCI domains of Cas9 are activated and induce a DNA double-strand break, with the HNH cutting the complementary DNA strand while the RuvCI cleaves the noncomplementary strand^3^. The double-strand break is repaired by the cell’s endogenous machinery, leading to insertions and deletions^4,6^. Homologous recombination can also occur, provided that a template for repair is present^4,6^.

Due to its high cutting efficiency in mammalian cells, the Cas9 enzyme from *Streptococcus pyogenes* (Sp) is the most widely studied of the Cas9 orthologues^4,7^, and has been used in the clinic^8^. SpCas9 is most efficient when the PAM sequence is composed of 5’-NGG (with N being any nucleotide), but also tolerates 5’-NAGs and 5’-NTGs^9,10^. This ability to accommodate mismatches between the target sequence and the sgRNA means that some off-target sequences are more accessible and unwanted mutations can occur^11,12^. This has led to a flurry of studies aiming to modify Cas9 to make it more specific or to change the compatibility of its PAM sequence. These studies used a variety of methods, including structure-based rational design^13–18^ and screening^19–21^. Luan et al^22^, for example, used molecular dynamics simulations and free energy perturbation to shorten the PAM recognition sequence of *Staphylococcus aureus* (Sa) Cas9. They found that free energy perturbation correlated well with gene editing efficiency in human cells. These results suggest that molecular dynamics simulations can assist the design of Cas9 variants quickly and cheaply.

Here we tested whether a similar approach could be used to design larger hybrids with extensive interacting interfaces. Indeed, whilst several variants with a single or a few amino acid changes have been used on the SpCas9 to change in the PAM preference of Cas9^17–19^ or improve on-target accuracy^13–16,20,21^, it has not yet been possible to reduce the size of the Cas9 enzyme. This is desirable because, SpCas9, with its 1368 amino acids, is too large to fit into an adeno-associated virus (AAV) together with its sgRNA for *in vivo* delivery. AAVs are one of best delivery systems because they display low immunoreactivity, have a wide tropism, and their viral DNA does not integrate easily within the host genome^23^. The size of SpCas9 means that two AAVs are required, one encoding the Cas9 enzyme, the other the sgRNA, resulting in increased cost, reduced efficiency, and an enhanced risk of an adverse immune reaction. To get around this problem, smaller Cas9 orthologues, like SaCas9 or *Campylobacter Jejuni* (Cj)Cas9, have been used in animal studies as they are small enough to be packaged into a single AAV with their sgRNAs^24–26^. However, they are less efficient than the SpCas9 and use different PAM sequences. SaCas9 recognizes NNGRRT (where R is a purine) and CjCas9 recognizes NNNNACA^26,27^. This means that most sequences targeted by SpCas9 are inaccessible to these smaller orthologues.

To overcome these issues, we aimed to develop new compact Cas9 hybrids by replacing the PAM Interacting Domain (PID) of SaCas9 and CjCas9 with that of SpCas9. Because sgRNAs are species-specific^28^, we also had to develop new sgRNA hybrids and ensure they interacted with the Cas9 hybrids protein. We used a two-pronged approach, taking advantage of *in silico* modeling to select four sgRNA/Cas9 pairs to test in human cells. We found that small point mutations or the presence of the sgRNA can drastically change the binding energy and the stability of the system, respectively. However, we found that the *in silico* approach did not predict the editing efficiency of the designed Cas9/sgRNA hybrid pairs in human cells, suggesting that molecular modelling and binding energy may not be sufficient to predict the function of large chimeric systems.

## Materials and methods

### Data and software Availability

Molecular dynamics simulations datasets are available here: https://github.com/DionLab/Cas9-hybrids, and as a version controlled here: 10.5281/zenodo.10029983.

### In silico models

The crystal structures required to run molecular dynamics were obtained from https://www.rcsb.org. PDB ID:4OO8 was used for SpCas9 in complex with its sgRNA and target DNA, this structure is at a 2.5Å resolution^5^. PDB ID:5CZZ was used for SaCas9 in complex with its sgRNA and target DNA this structure is at a 2.6Å resolution^27^. We note that the SaCas9 structure lacked stem loop 2 of the sgRNA. PDB:5X2G was used for the CjCas9 in complex with its sgRNA and a target DNA which was available at a 2.4Å resolution^29^. In this structure, the CjCas9 protein lacked the HNH domain, which was replaced with a GGGSGG linker. The SaSpCas9 hybrid is the hybrid between the N-terminal of SaCas9 (1-910AA) and the PID of SpCas9 (1100-1368AA). Similarly, the CjSp Cas9 hybrid is a fusion between the N-terminal of the CjCas9 (1-828AA) and the same Sp PID. To make the nickase mutations, we used the D10A mutation for the SaSp Cas9 and the D8A for the CjSpCas9 (Supplementary Table 1), and created the change using the protein editing function in Molecular Operating Environment (MOE) 2019. Cas9 hybrids and sgRNA hybrids (Supplementary Table 1 and 2) were made using MOE 2019 and PyMOL v2.5.4. An energy minimization was run on MOE to avoid any clashes between the added PID and rest of the protein prior to molecular dynamics simulations. We also designed the SaSp protein using ColabFold v1.5.3. Alphafold2_multimer_v3 was used for complex prediction. A total of 3 recycles count (default) were used to predict the model. A total of 200 max iterations were used with a greedy pairing strategy. This approach, however, did not yield a useable model. Alphafold2 is currently unable to co-model with DNA or RNA structures and as such the model produced contained segments of protein which were positioned in such a way that the DNA/RNA would not be able to fit in the protein anymore. Further to this, Alphafold2 works in such a way that it is good at predicting proteins based on an evolutionary model, our hybrids are synthetic structures and not a natural protein, Alphafold2 is therefore not designed to model such a protein and we chose instead to use the MOE software to design our structure being careful to preserve important structural features of both protein and DNA/RNA.

### Molecular dynamics simulation

All the preparation steps and analysis for molecular dynamics simulation were carried out using GROMACS version 2018.2^30^. The forcefield AMBER03 was employed to create a stable environment for the system^31^. The protein-RNA-DNA complex was set in a 0.9 nm box filled with TIP3 water molecules. The system was neutralized either by adding Na+ or Cl-ions depending on the overall charge. An integration time step of 0.001 ps was used. The complex was energy minimized to find the best position for the protein to avoid any clashes between the different structures. The energy minimization was run for a maximum of 3000 steps and a maximum energy barrier of 1000 kJ/mol/nm. During the molecular dynamics simulation, the v-rescale coupling method regulated the temperature to 310 K. A Berendsen barostat was coupled to the system to regulate the pressure with a reference of 1 bar. The whole simulation was composed of 5 different groups: protein, ions, solution, DNA, and RNA. The molecular dynamics simulation was run for a total of 100 ns.

### Stability data analysis

The stability of every complex was studied by analyzing the Root Mean Square Deviation (RMSD) and the Root Mean Square Fluctuation (RMSF). Each of these are modules built-in GROMACS. The RMSD was extracted for every structure of the complex (protein-RNA-DNA) using the rms command and given every 10 ps. An average of the RMSD was made only including data from 10 to 100 ns as the first nanoseconds serve to warm up the system. The RMSF was extracted for every residue of every structure present in the complex using the rmsf command.

### Binding energy

The Molecular Mechanics Poisson-Boltzmann surface area (MMPBSA) method was used to precisely determine the interaction between each base of the sgRNA hybrid and the protein hybrid. Recently, a GROMACS integrated tool was developed to perform these calculations^32^. The files and instructions needed to run it were found on: https://rashmikumari.github.io/g_mmpbsa/. This method combines calculations of potential energy in a vacuum, polar solvation energy, and non-polar solvation energy. The binding energy with g_MMPBSA, even if less computationally hungry than other methods, remains time-consuming. To reduce the computational burden and make the analysis possible, the last 100 frames of the molecular dynamics simulations were extracted and used to calculate the binding energy between the sgRNA and the protein.

### Cell lines plasmids and culture conditions

HEK293-derived GFP(CAG)_0_ and GFP(CAG)_101_ cells were a gift from John H. Wilson^33^. They were maintained at 37°C with 5% CO_2_ in DMEM supplemented with 10% FBS, 100 U ml^-1^ penicillin, 100 μg ml^-1^ streptomycin, 15 μg ml^-1^ blasticidin, 150 μg ml^-1^ hygromycin, and, during the experiments, 1 µg ml^-1^ of doxycycline diluted in water. Both lines were regularly tested for mycoplasma using a service from Eurofins. The cells remained mycoplasma-free throughout the experiments. They were also genotyped using the Mycrosynth AG service and were determined to be HEK293.2sus as previously determined^34^.

Transfections were done as before^35^ by seeding 400,000 cells in a 12-well plate well on day 0 and transfecting 1 µg of plasmid using lipofectamine 2000. GFP intensities were measured using an Attune NxT flow cytometer and analyzed using the FlowJo software version 10.8.1. The plasmids used in this study (Supplementary table 3) will be available via Addgene shortly.

For the GFP editing assays, we used GFP(CAG)_0_ cells along with 4 well-characterized sgRNAs against GFP^36^ (Supplementary Table 4). The GFP levels in the cells were measured at day 6, 8, and 10 post-transfection. The flow cytometry data were analyzed using FlowJo v.10.8.1

The results were confirmed using a T7 endonuclease I assay using GFP(CAG)_0_ cells transfected with GFP sgRNAs target 1 or 3 (Supplementary Table 4). The targeted regions were amplified using the primers oVIN-3474 and oVIN-3475 for sgRNA 1, 3 and 4 (Supplementary Table 5). The target region for sgRNA 2 was amplified using oVIN-3476 and oVIN-3477 (Supplementary Table 5). The PCR protocol used was 95°C for 5min, followed by 35 cycles at 95°C for 30’’, 55°C for 30’’ and 72°C for 1 min followed by a final step at 72°C for 10 min. The annealing of PCR products was performed by mixing 200 ng of DNA in 1x NEBuffer2. After annealing, 10 U of T7 endonuclease I (NEB) was added and incubated for 15 min at 37°C. For repeat instability assays using the GFP reporter, GFP(CAG)_101_ cells were cultured as above, but with dialyzed FBS (Merck). The cells were transfected on days 0, 4, and 8 before being analyzed for GFP intensity by flow cytometry on day 12. The resulting datasets were analyzed using FlowJo v.10.8.1 The plasmids used for transfection can be found in Supplementary Table 2.

### Protein quantification and western blot

Proteins were extracted using commercial RIPA buffer (Fisher Scientific) and quantified using Pierce BCA protein assay (ThermoFisher Scientific). The proteins were separated on a 4-12% Bis tris gel (ThermoFisher Scientific) and transferred onto a nitrocellulose membrane (Bio-Rad). The membrane was blocked for 1 h in 5% milk in PBS-T then incubated overnight with an anti-flag antibody (Sigma) or actin antibody (Sigma) at 4°C. The secondary blotting was performed after 1 h incubation in an Alexa Fluor 680 anti-mouse antibody (Invitrogen). The blot was imaged using Licor Odyssey CLX.

### Statistics

Two-tailed Mann-Whitney U tests were performed to determine statistical significance between the RMSD values of the hybrids and their relevant wild type Cas9 orthologues and between the two CjSpCas9. We used a Kruskal-Wallis test to determine statistical significance between the RMSD values of the SaSp hybrids. We used Graphpad Prism (version 10.0.0) to calculate the p-values.

## Results

### Molecular dynamics and binding energy identify functional changes in Cas9

To test whether our *in silico* approach could identify meaningful functional changes, we ran molecular dynamics simulations of the SaCas9 complex with and without its sgRNA. We reasoned that the presence of the sgRNA would have a large impact on the dynamics of the Cas9 enzyme. RMSD and RMSF values were extracted from the simulations. RMSD is a measure of the average distance for a group of atoms between a reference structure and the resulting structure after a specified amount of time. The RMSF relates to the movement of each individual residue over the time of the simulation. The SaCas9 without its sgRNA reached a maximum RMSD of 1.22 nm but remained stable beyond 40 ns (Figure 1A), with an overall RMSD average of 1.1 nm. In contrast, the SaCas9 in complex with its sgRNA was more stable with an average RMSD of 0.33 nm, and a maximum of 0.37 nm (Figure 1A). Consistent with these data, the RMSF analysis showed that every domain of the protein is stabilized by the presence of its sgRNA (Figure 1B). The REC lobe, the bridge helix, RuvCI, and HNH domains, were the most impacted. We conclude that changes in RMSD and RMSF values reflect large scale changes in SaCas9.

**Figure 1:**
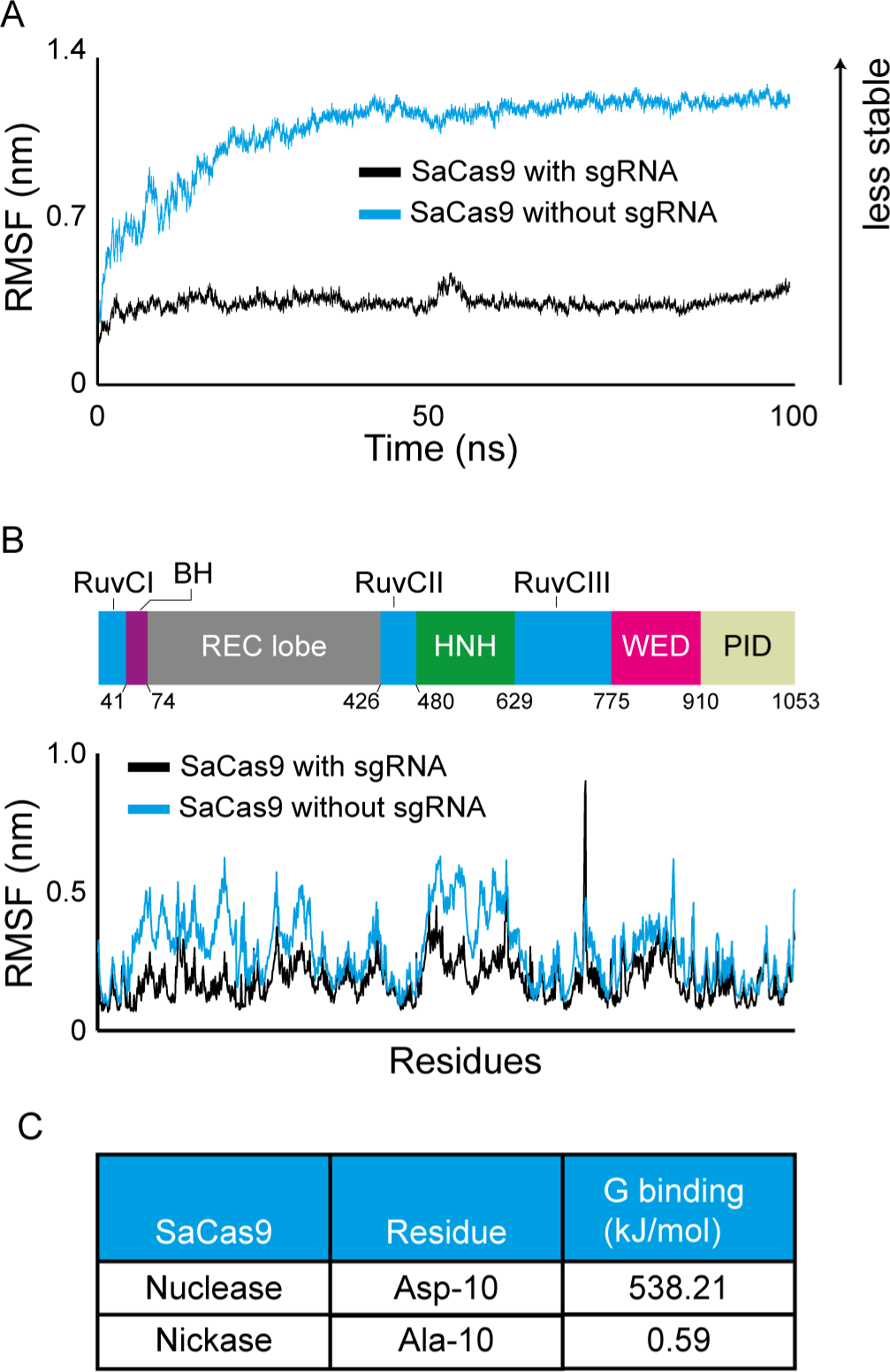
Molecular dynamics simulations and energy binding calculation identify functionally relevant changes in Cas9 activity. A) Root mean square deviation (RMSD) of SaCas9 with or without its sgRNA over 100 ns simulations. B) Root mean square fluctuation (RMSF) of SaCas9 with or without its sgRNA. (BH = bridge helix, WED = wedge domain, PID = PAM interacting domain). C) G binding values at position 10 of SaCas9 comparing the nuclease (D10) to the nickase (A10).

To determine whether we could find functional differences using binding energy measurements, we compared SaCas9 to the D10A mutant, which turns the nuclease into a nickase ^3^. To do so, we used the MMPBSA approach^32,37,38^, which works by calculating the overall energy for Cas9 in solution compared to that of Cas9 in a vacuum. The same is done for the sgRNA. From these data, we can calculate the energy difference between Cas9 and its sgRNA in solution as well as in a vacuum, which gives us an approximation of the binding affinity between the two molecules for each amino-acid (Supplementary Figure 1A). We reasoned that the binding energy for the mutated residue should change if this method is predictive of a change in function. Moreover, the rest of the protein should change very little, if at all, by this amino acid change. To test this directly, we calculated the binding energy associated with every residue extracted from the SaCas9 simulation and compared them to that of the SaCas9 D10A variant. We found that the G binding of D10 was 912 times greater than that of A10 (Figure 1C). This is in contrast to the other residues, which on average changed by 0.39 kJ/mol (Supplementary Figure 1B). Together, these analyses show that the molecular dynamics simulation together with the binding energy calculations robustly predicts functional changes.

### The Cas9 hybrid complexes are stable in silico

Next we tested whether we could use this computational approach to design multiple compact Cas9/sgRNA hybrid pairs to be tested in human cells. This was done in two steps. First, we performed molecular dynamics simulations and determined the RMSD and RMSF of the hybrid pairs and then determined which ones to take forward using energy binding calculations. We initially designed 9 such pairs *in silico*, 7 SaSp pairs and 2 CjSp hybrid pairs (Figure 2A-B). For all these, as well as for the wild type versions of SpCas9, SaCas9, and CjCas9, we performed molecular dynamics simulations and analyzed the stability of these hybrids (Figure 2A-J and Supplementary figure 2A-F and supplementary3A-B). We found that in the presence of a sgRNA the RMSD for the SaSpCas9 or CjSpCas9, regardless of the sgRNA hybrid used, was systematically higher than the SaCas9, SpCas9 or CjCas9 coupled to their wild type sgRNAs (Figure 2C,D – P value < 0.0001). However, for both hybrids, a plateau was reached, showing that the proteins arrived at a stable conformation.

**Figure 2:**
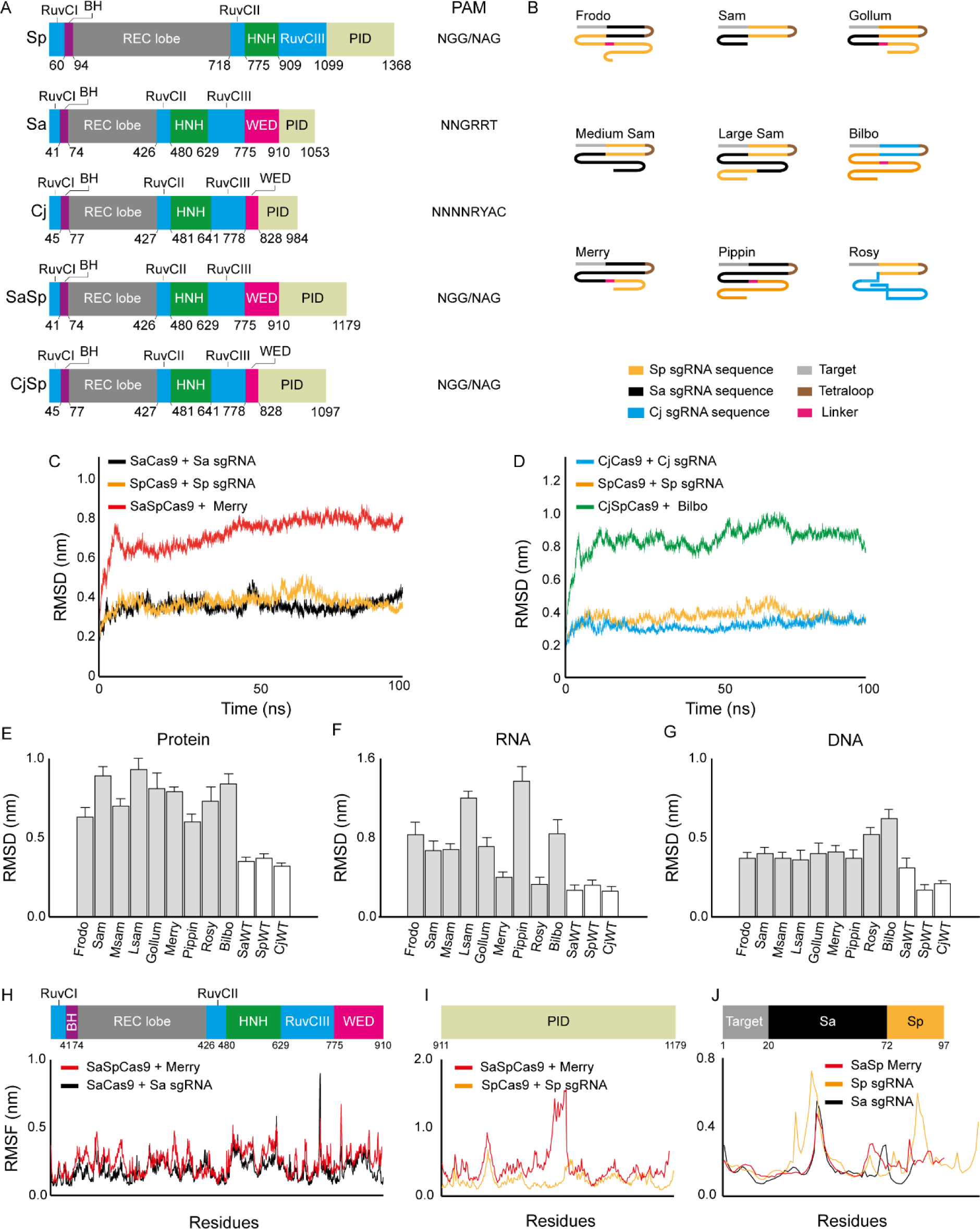
Molecular dynamics simulations of Cas9 hybrids. A) Domain organization of the Sp, Sa, and Cj Cas9 as well as the hybrids tested here. The PAM sequence is either the reported ones or, for the hybrids, the expected PAM. The numbers indicated the amino acids at domain boundaries (BH = bridge helix, WED = wedge domain, PID = PAM interacting domain). B) Schematic representation of the sgRNA hybrids tested here. C) RMSD of SpCas9, SaCas9, and SaSpCas9 with their specific sgRNAs during 100 ns. The SaSp sgRNA presented is Merry. D) RMSD of SpCas9, CjCas9, and CjSpCas9 with their specific sgRNAs over a simulation of 100 ns. The CjSp sgRNA presented is Bilbo. E-G) Average RMSD for the protein (E), sgRNA (F), and DNA (G) in all 12 systems modeled. H) RMSF of SaCas9 with its sgRNA (black) and SaSp Cas9 in combination with Merry (red) for the N-terminal common to both proteins. I) RMSF of the PID of SpCas9 with its sgRNA (yellow) and SaSpCas9 in combination with Merry (red). J) RMSF of SpCas9 sgRNA (yellow), SaCas9 sgRNA (black) and SaSpCas9 sgRNA (red).

When comparing the protein RMSD for the 7 SaSp models, we found that Large Sam (LSam) showed the highest maximum average at 0.93 nm whereas Pippin had the lowest maximum average at 0.60 nm (Figure 2E). For the CjSp, Rosy had the lowest value with 0.73 nm. The same protein hybrids showed different RMSD values and were significantly different from one to another (P<0.0001) presumably, because of how they interacted with their respective sgRNAs. Analysis of the RNA RMSD showed that Pippin had the highest value with 1.37 nm whereas the lowest values were seen with Merry (0.40 nm, Figure 2F). For the CjSp hybrids, again, Rosy had the lowest value with 0.33 nm. The DNA RMSD did vary significantly between the SaSp systems (P<0.0001) and ranged between 0.36 nm (Msam) and 0.41 nm (Merry; Figure 2G). For the CjSp, Rosy had the lowest value of 0.52 nm. Altogether, the RMSD data show that the hybrids had consistently higher values than the three wild types, yet they were below what we observed for SaCas9 without the presence of a sgRNA (Figure 1).

To complement these findings, we determined the RMSF values of each residue in the proteins and sgRNAs for each hybrid pair. We found that the N-terminal of the SaSpCas9 coupled to Merry (Figure 2H) and to Gollum (Supplementary Figure 2A) matched closely the wild type SaCas9 coupled to its sgRNA, with the HNH domain being the most fluctuating domain for both proteins. The PID of the SaSp hybrids (Figure 2I and Supplementary Figure 2A-F) were more mobile than the Sp wild type pair, especially for some atoms around residue 1000. This may be explained by the energy minimization step that moved the initial position of these residues away from other atoms, giving it more space to move. The RMSF for SaSp Merry sgRNA was closest to that of the Sp and Sa sgRNAs (Figure 2J) with the exception of stem loop 2 from Sp wild type showing more fluctuations. Similar to the SaSpCas9 hybrids, we found that the CjSpCas9 hybrids were consistently more mobile than the wild types (Supplementary Figure 3A-B). Altogether, the RMSD and RMSF data suggest that the Cas9 hybrids that we designed are stable during molecular dynamics simulations.

### Binding energy of the Cas9/sgRNA hybrids closely match that of wild type complexes

To determine which hybrid to test in cellular systems, we analyzed the binding energy of every SaSp hybrid pair. The analysis of the SaCas9 complex could readily identify nearly all residues previously shown to interact with the sgRNA in the crystal structure^27^, as well as additional ones (Figure 3A and Supplementary Figure 4). This added confidence that the residues with the smallest binding energy were indeed structurally important. Comparing the binding energy in the residues interacting with the sgRNA, we found that some SaSp sgRNAs, especially, Merry and Gollum, showed very similar binding energy compared to the SaCas9 and its sgRNA (Figure 3A and Supplementary Figure 4A). By contrast, Frodo showed more extreme values (Figure 3B). Similarly, the CjSp complexes Rosy and Bilbo showed similar values as the CjCas9 complex, but Rosy had one amino acid that interacted with the sgRNA in CjCas9, Arg-63, which lost its binding affinity (Supplementary Figure 5B). We conclude from these binding energy calculations that the best designs, as defined as those that most closely resemble the structural characteristics of SaCas9, SpCas9, and CjCas9, were Merry and Gollum for the SaSpCas9 hybrid and Bilbo for the CjSpCas9 hybrid. We also tested Frodo in cells as an example of a less optimal design.

**Figure 3:**
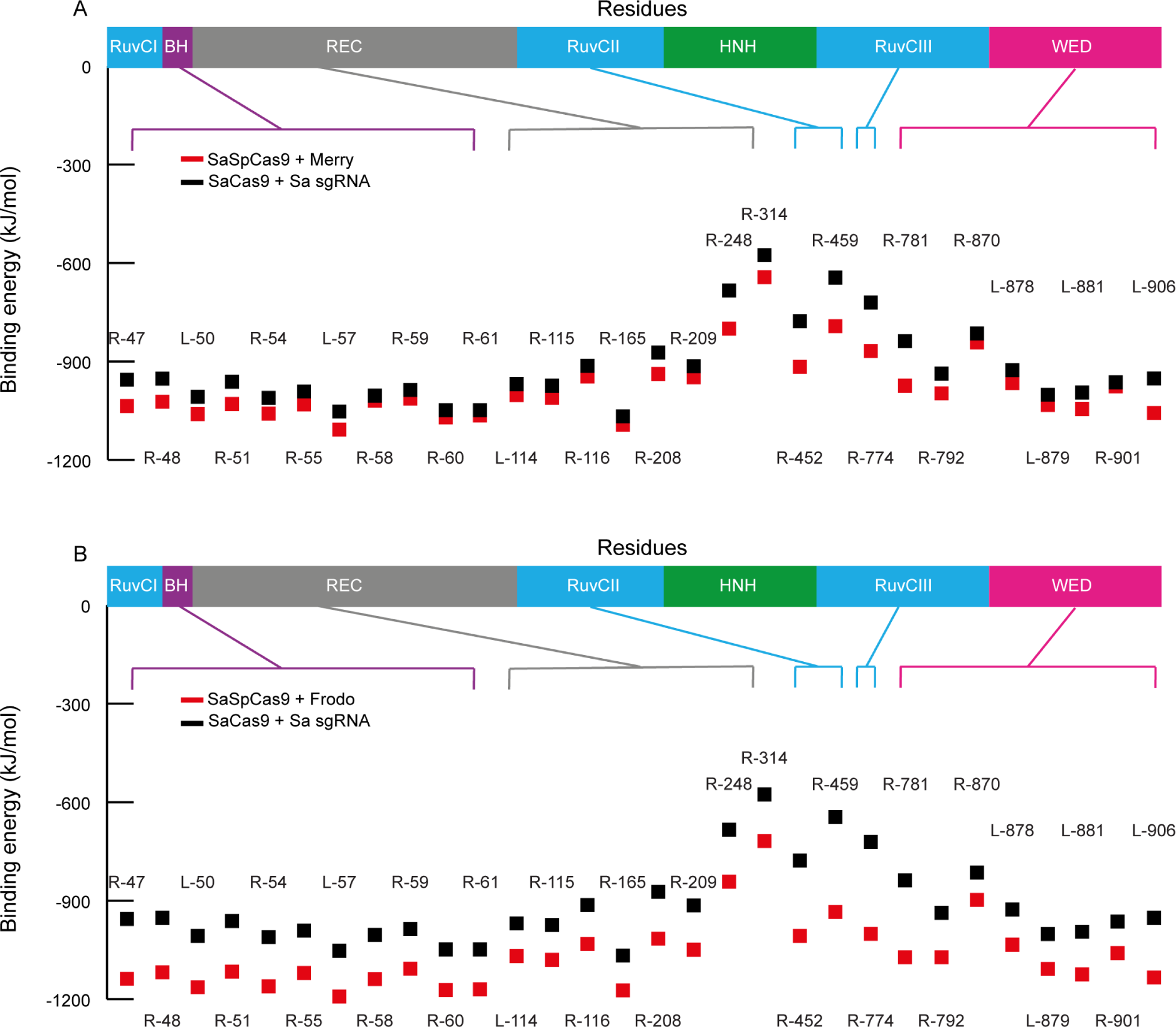
Binding energy calculations for SaSp hybrids. A-B) Binding energy for each amino acid known to interact with the sgRNA for SaCas9 with its sgRNA (black) versus SaSpCas9 with Merry (A) or Frodo (B) (red). BH = bridge helix, WED = wedge domain, PID = PAM interacting domain.

### The Cas9 hybrids failed to edit DNA in human cells

Next we wondered whether the designed Cas9 hybrids could edit DNA in cells. We cloned our SaSpCas9 together with either Merry, Gollum, or Frodo. We transfected them into GFP(CAG)_0_ cells and showed that they all expressed to comparable levels (Supplementary Figure 6A,B). Then, we cloned each sgRNA hybrid with four different target sequences against the GFP gene (Figure 4A and Supplementary table 4). This is convenient because gene editing produces insertions and deletions that inactivate GFP. We could then readily detect the drop in GFP intensity using flow cytometry. Using this approach, we confirmed that the SpCas9 was active, leading to a loss of GFP intensity of up to 15% for three of the four target sequences (Figure 4C). By contrast, we found that the SaSpCas9/sgRNA hybrid pairs changed GFP expression very little, if at all, suggesting that they were not functional.

**Figure 4:**
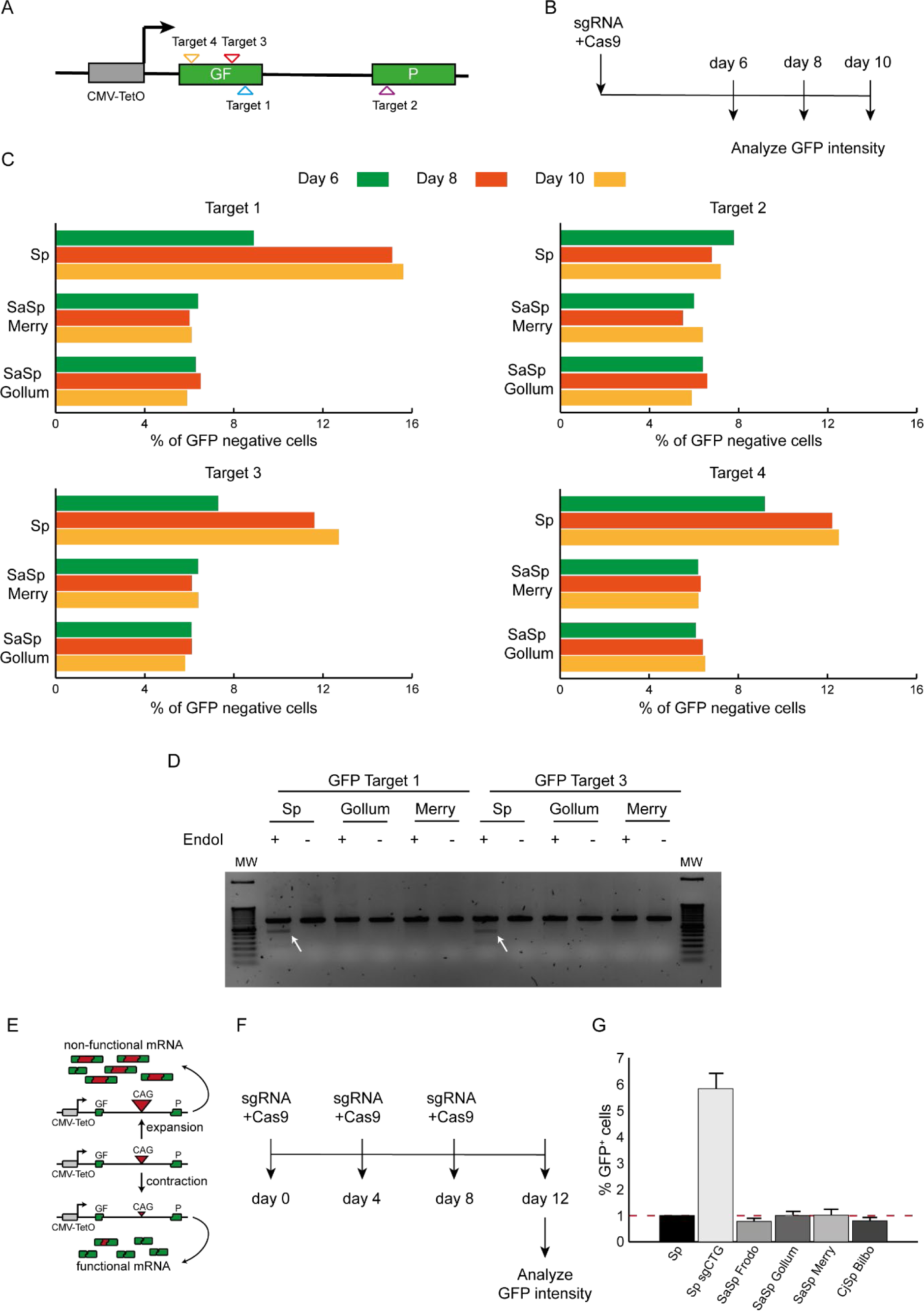
The Cas9 hybrids fail to edit DNA in cells. A) Schematic representation of the GFP gene construction used for the editing assay. Each position targeted by the sgRNA used are shown with different triangles. This representation is not to scale. CMV-TetON is a doxycycline-inducible promoter. B) Timeline of experiment used for the GFP assay. C) Percentage of GFP negative cells for the 4 targets used at day 6 (green), 8 (orange) and 10 (yellow) for Sp, SaSp Merry and SaSp Gollum. D) T7 endo I assay results for targets 1 and 3. The white arrows indicate the cleavage products indicative of efficient editing. E) GFP-based assay used to detect contraction of the CAG repeats length. The repeat interferes with the splicing of the reporter in a size-dependent manner. F) Timeline experiment for the GFP-based assay to detect CAG contractions. G) Percentage of GFP positive cells for each condition. Dashed lines represent the gate containing the brightest 1% of the cells transfected with Cas9 only. Previous work^40^ has shown that this population is enriched in shorter repeats.

The GFP assay is an indirect measure of gene editing. To complement it, we used a second assay, the T7 endonuclease I assay, which detects edits at the DNA level. The insertions and deletions, once amplified and hybridized to unedited strands, form mismatches. These mismatches are then cut by the T7 endonuclease I, producing smaller DNA fragments that can be resolved on an agarose gel. We extracted DNA from the GFP assay at day 10 and performed the T7 endonuclease I assay for targets 1 and 3 as they showed the best results in editing the GFP gene. The results confirmed that whereas mismatches were generated upon SpCas9 transfection, the SaSpCas9 with Merry or Gollum induced no detectable edits (Figure 4D), suggesting they are not functional.

As an independent method to confirm the lack of activity of the hybrids in cells, we used a well characterized assay for the contraction of CAG repeats in GFP(CAG)_101_ ^33^. The assay works because CAG repeats interfere with the splicing of the GFP reporter in a size-dependent manner with longer repeats producing less GFP (Figure 4E). The SpCas9 nickase (D10A mutant) together with a sgRNA that targets the repeat tract itself induces contractions of the CAG/CTG repeat, thereby increasing GFP levels in these cells^39,40^. We therefore transfected SaSpCas9 together with Frodo, Merry, or Gollum, as well as CjSpCas9 with Bilbo and measured GFP levels (Figure 4F). As expected, the SpCas9 enzyme increased GFP intensities, indicating contraction of the CAG repeat, whereas the Cas9 hybrids did not (Figure 4G). Together, these results show that the SaSp and CjSp hybrid pairs are not functional in human cells.

## Discussion

In this study, we tested the hypothesis that we could use molecular dynamics simulation and binding energy data to design *in silico* Cas9 hybrids that retained the PAM properties of SpCas9, but were small enough to fit within a single AAV. In addition, we found that the binding energy of the hybrids closely mirrored that of wild type Cas9 enzymes, thereby predicting edits in cells. This was not the case. Most other variants of SpCas9 and SaCas9 in the literature differ from the wild type by just a few amino acids and were designed to decrease the frequency of binding off-targets or to change their PAM recognition^13–21^. Similarly, three chimeric Cas9 enzymes have been created that change PAM sequence requirements^41,42^. At first glance these studies appear similar to what we have done here in that the PID of one orthologue was fused to the N-terminal of another. However, the fusions from those previous studies were between SpCas9 N-terminal and the PID of either *Streptococcus macacae* or *Streptococcus thermophilus* Cas9. Both of these orthologues are closely related to SpCas9 and their sgRNAs are interchangeable. Moreover, these hybrids do not reduce the overall size of the resulting proteins, which was our primary goal. Our results suggest that although powerful enough to predict the functional impact of one or a few residues, molecular dynamics simulation and binding energy values are not sufficient for predicting function when the changes to the systems are on a larger scale.

It might be possible to further optimize the hybrid pairs by changing specific amino acids while minimizing the RMSD and RMSF. This may help improve the Cas9/sgRNA interactions, stabilize the hybrids further, and lead to a functional outcome. But this approach is predicated on the hypothesis that the parameters are predictive of function, which we have no evidence for here. It is also possible that the sgRNA hybrids were not expressed well enough to lead to robust gene editing.

Alternatively, the approach was perhaps successful, but the PAM requirements from the SpCas9 were not retained. Indeed, some SpCas9 with PAM variants are due to changes lying outside the PID^18,19^. If this was the case, then the target sequences we used here would not be recognized by our hybrids, explaining the lack of edits.

Further work will be required to make small Cas9 variants suitable for *in vivo* work that retain the SpCas9 PAM interacting properties. This may include taking the Cas9 hybrids and screening empirically for sgRNAs hybrids that bind to the enzyme or adapting the PAM recognition of smaller orthologues. Improving the structure-function prediction of Cas9 variants would benefit a wide range of applications.

## Acknowledgements

We thank members of the Dion group for critical reading of the manuscript. This research was undertaken using the supercomputing facilities at Cardiff University operated by Advanced Research Computing at Cardiff (ARCCA) on behalf of the Cardiff Supercomputing Facility and the HPC Wales and Supercomputing Wales (SCW) projects. We acknowledge the support of the latter, which is part-funded by the European Regional Development Fund (ERDF) via the Welsh Government.

## Author contributions

AM performed all the experiments and designed them with the help of VD (for cell work) and GM (for *in silico* work). AM generated all figures and the manuscript was written in collaboration between all three authors.

## Supplementary Materials

**Supplementary figure 1:**
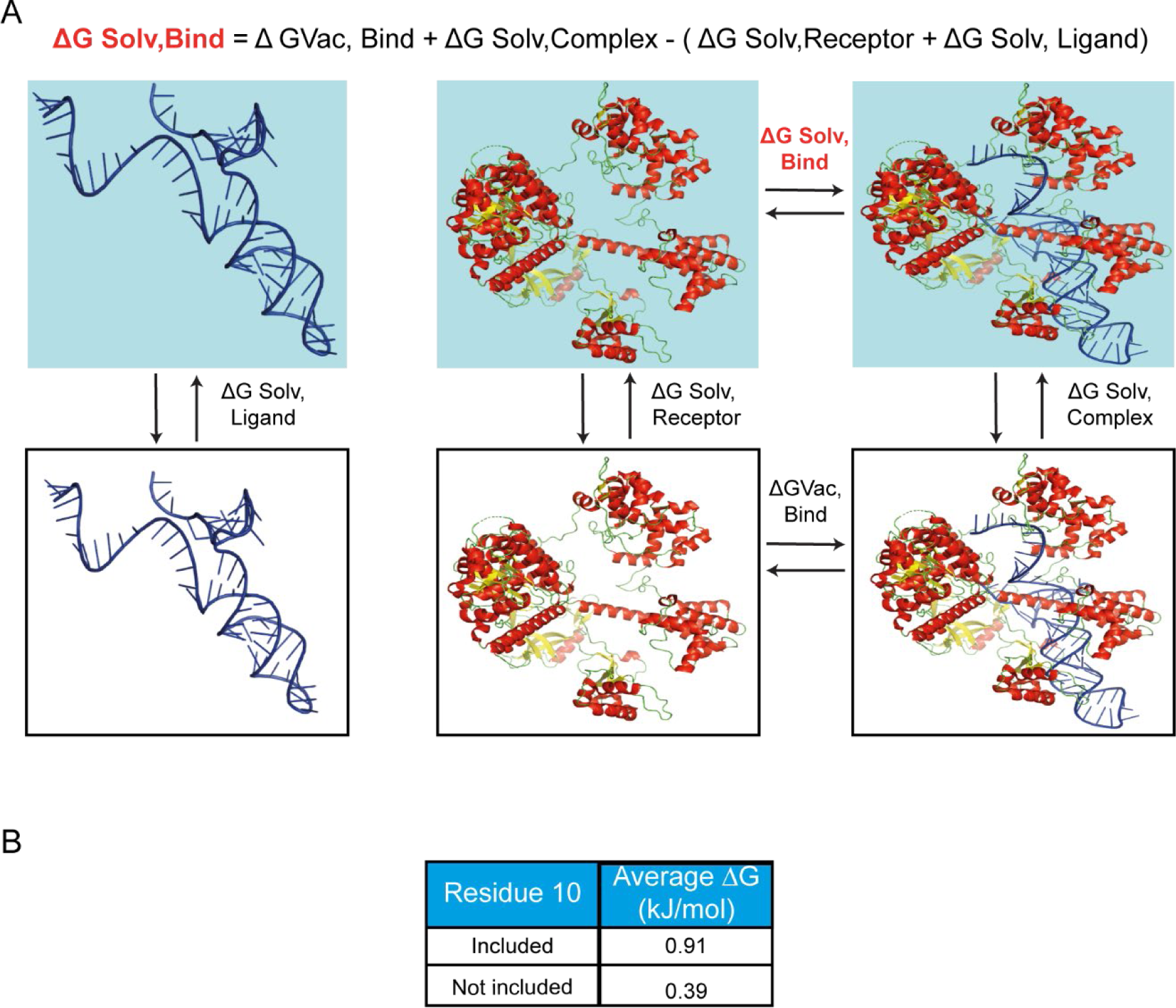
Binding energy calculations between the Cas9 protein and its sgRNA. A) Principle behind binding energy calculations. The blue boxes represent the system in water whereas the white boxes are in vacuum. B) An average of the ΔG binding for each residue between the nickase and nuclease. ‘Included’ means that the average ΔG was calculated for every residue in the Cas9 protein, whereas ‘not included’ excludes the amino acid that changes between the nuclease (D10) and the nickase (A10). This shows an outsize contribution of this residue to the overall binding energy of the protein.

**Supplementary figure 2:**
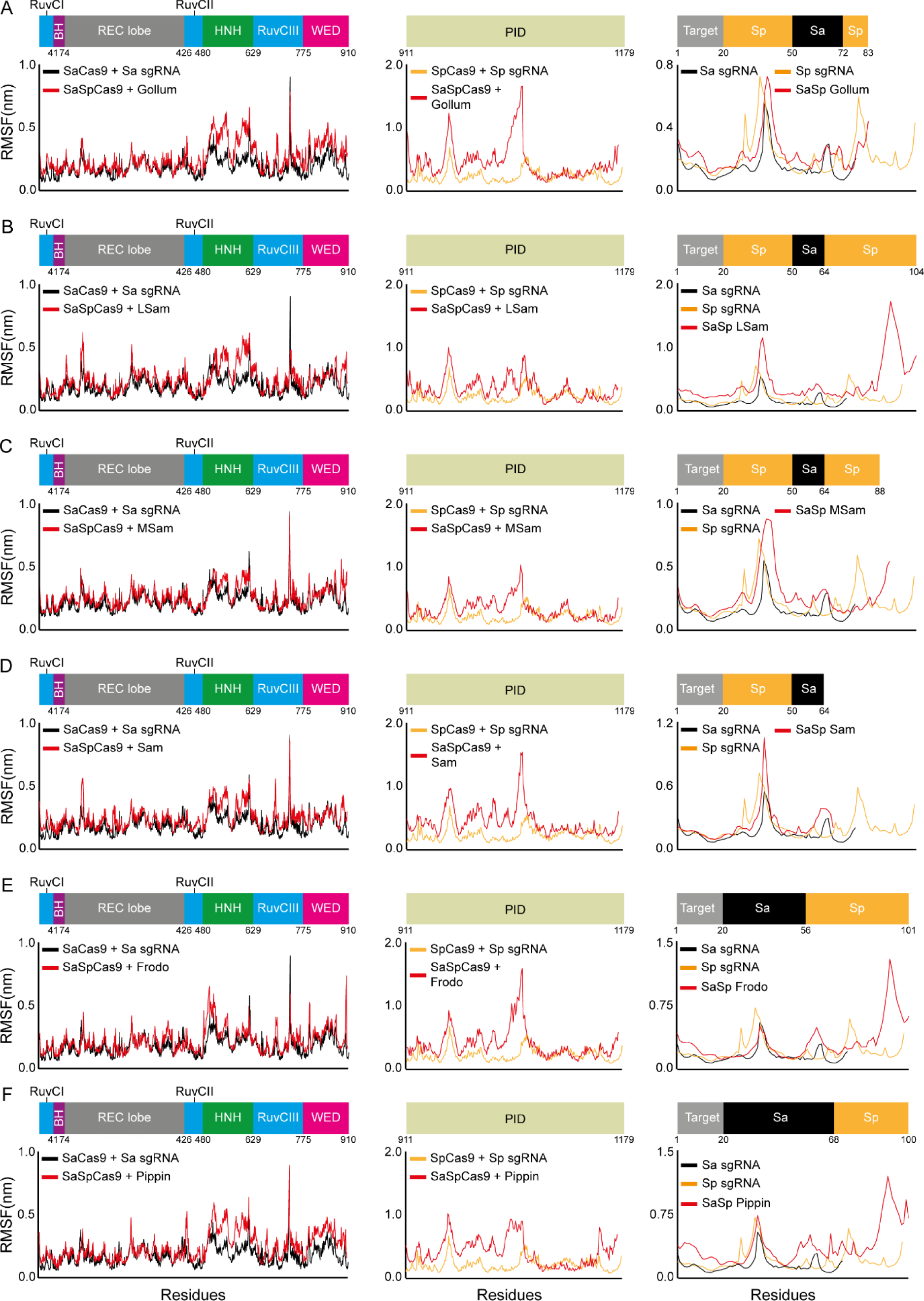
RMSF analysis from molecular dynamics simulations for the SaSp hybrids: A - F) RMSF of SaSp hybrids (red), SaCas9 (black), and SpCas9 (orange) together with their cognate sgRNAs. A) Gollum, B) LSam, C) MSam, D) Sam, E) Frodo, F) Pippin. Left: N-terminal, middle: PID, right sgRNA. This figure shows that all the hybrid pairs are expected to be stable.

**Supplementary figure 3:**
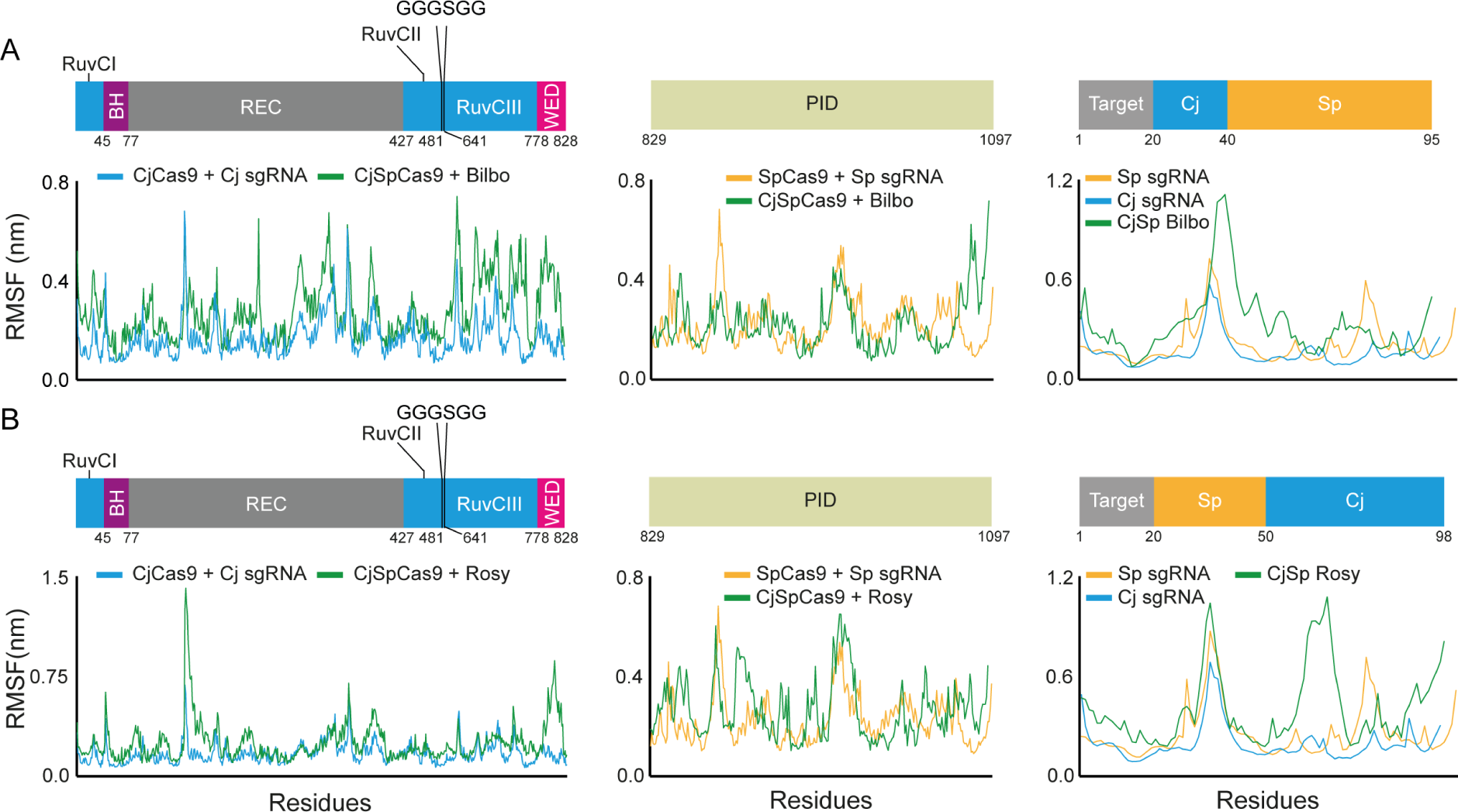
RMSF analysis from molecular dynamics simulations for the CjSp hybrids: A - B) RMSF of CjSp hybrids (green), CjCas9 (blue) and SpCas9 (orange), together with their cognate sgRNAs. A) Bilbo, B) Rosy. Left: N-terminal, middle: PID, right sgRNA. This figure shows that all the hybrid pairs are expected to be stable.

**Supplementary figure 4:**
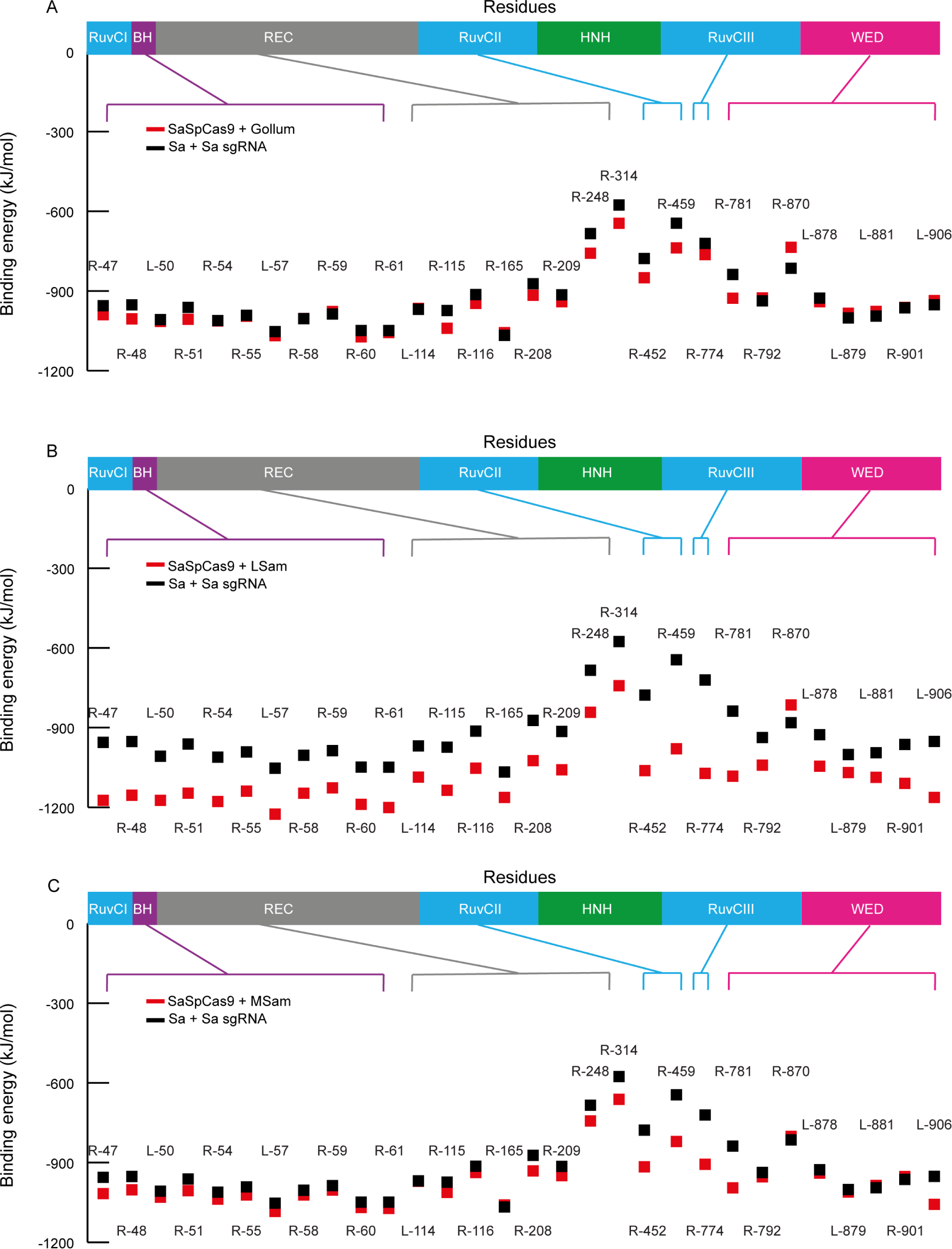
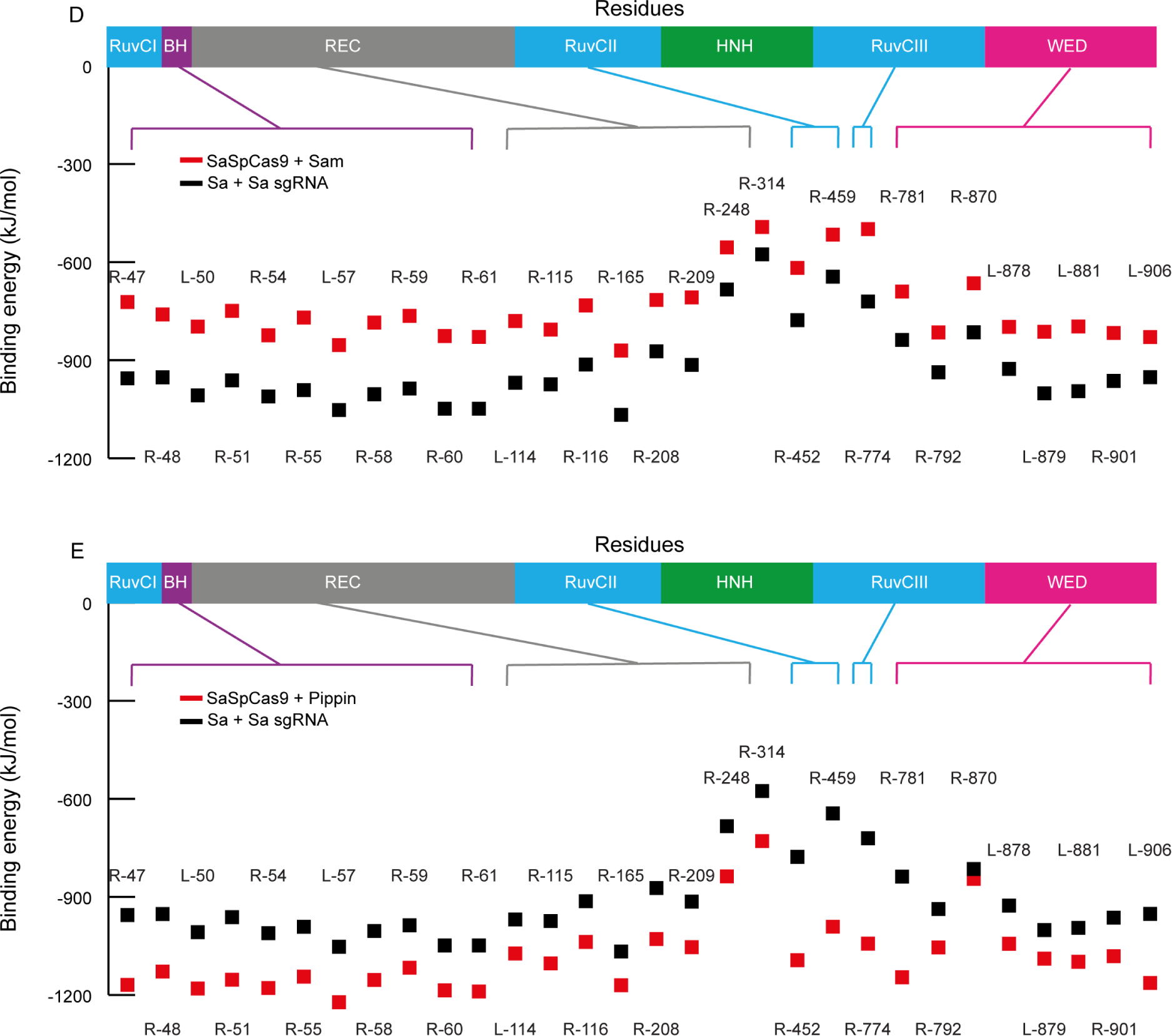
Binding energy calculation for the SaSpCas9 hybrids. The binding energy for each amino acid known to interact with the sgRNA from the crystal structures are shown for comparing A) SaCas9 (black) to SaSpCas9 (red). A) Gollum, B) LSam, C) MSam, D) Pippin, E) Sam. This figures shows that the sgRNAs are expected to make the contacts with the sgRNAs as predicted from the structural information available.

**Supplementary figure 5:**
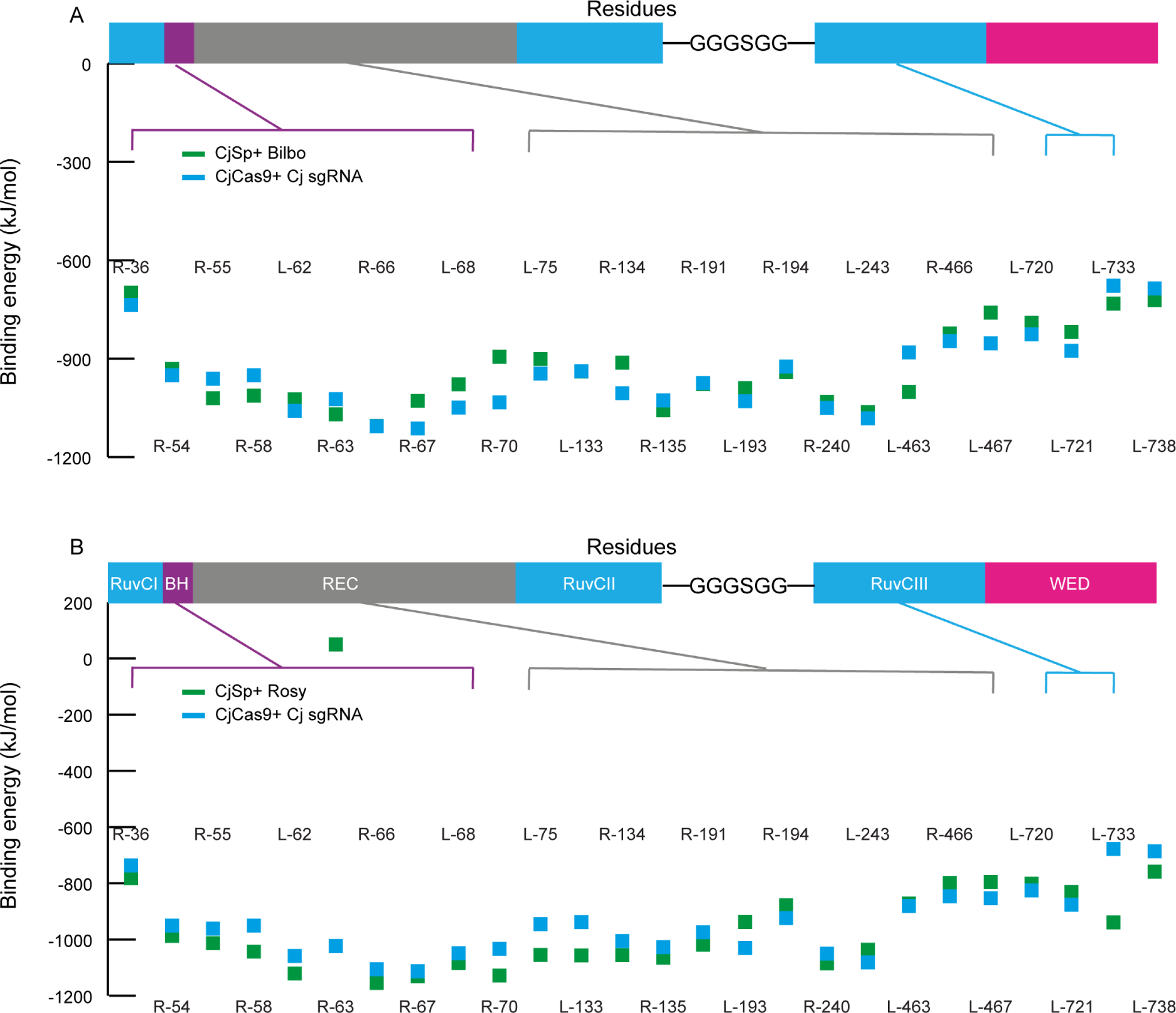
Binding energy calculation for the CjSPCas9 hybrids. The binding energy for each amino acid known to interact with the sgRNA from the crystal structures are shown for comparing CjCas9 (blue) to CjSpCas9 (green). A) Bilbo, B) Rosy. This figures shows that the sgRNAs are expected to make the contacts with the sgRNAs as predicted from the structural information available.

**Supplementary figure 6:**
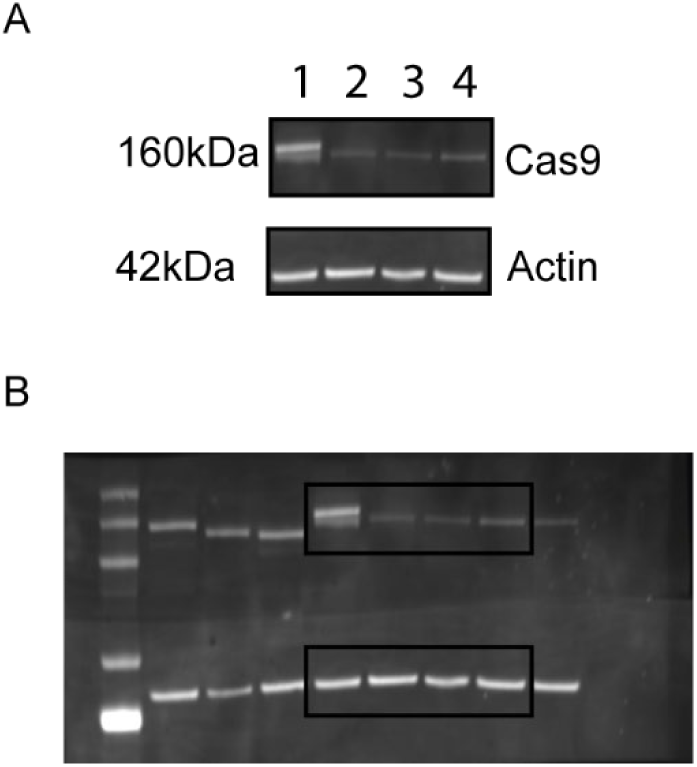
Expression of Cas9 in HEK cells: A) Expression of 1) SpCas9 together with its sgRNA, 2) SpCas9 with Gollum, 3) SaSpCas9 plus Frodo, and 4) SaSpCas9 alongside Merry. B) Uncropped western blot for Cas9 expression. Black boxes indicate where the blots were cropped. Note that the membrane was cut prior to blotting but imaged together.

**Supplementary Table S1:**
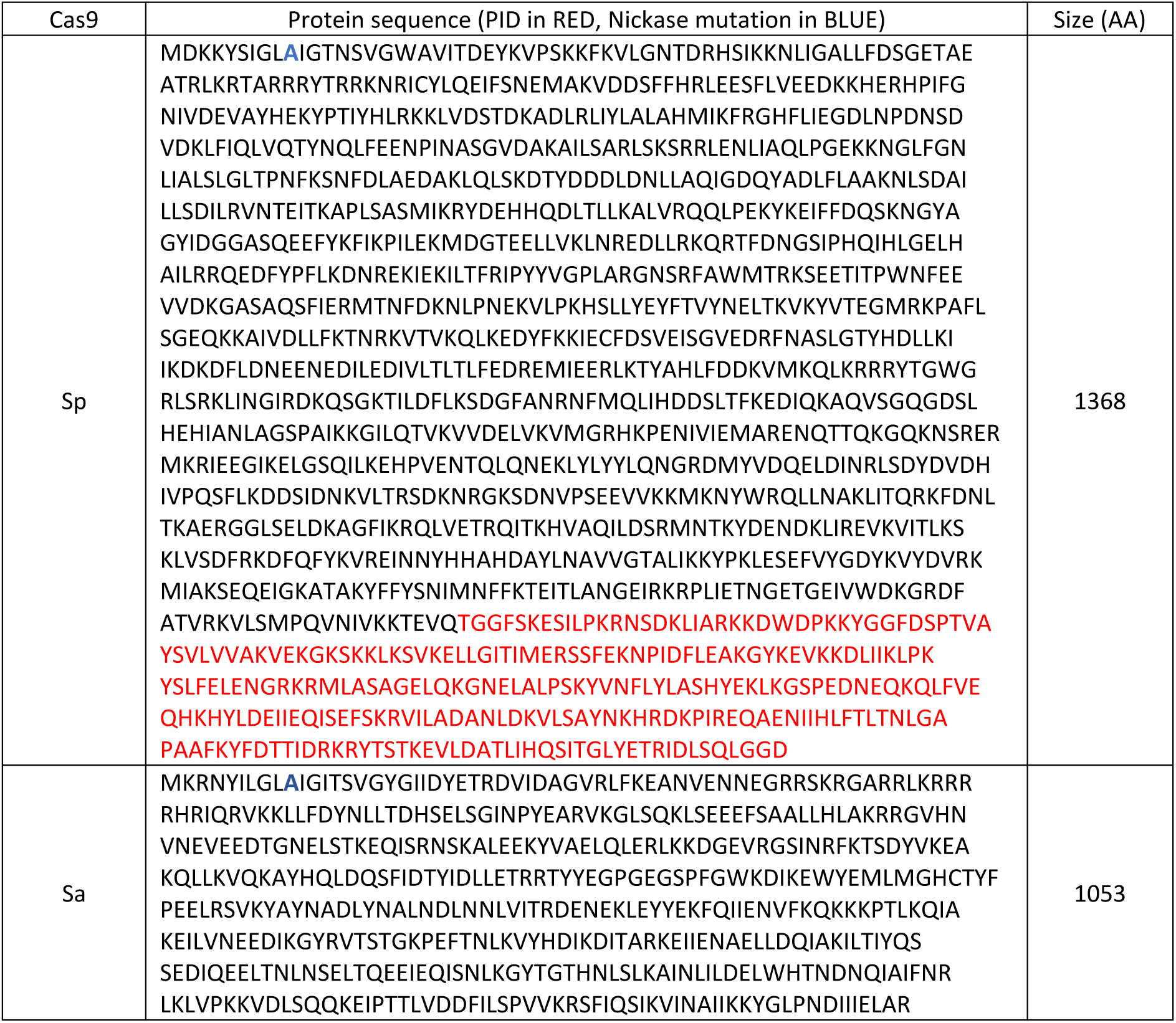
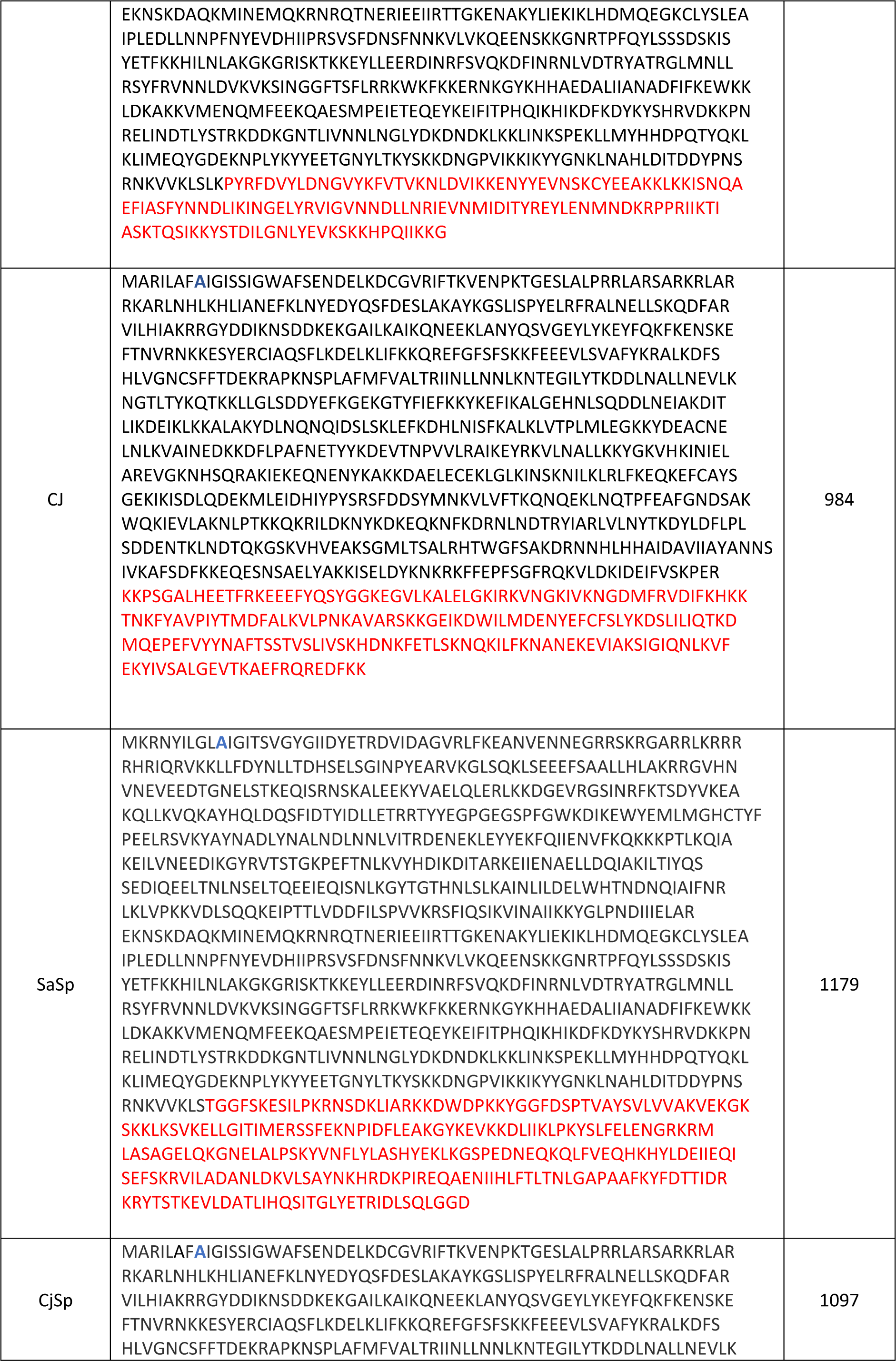
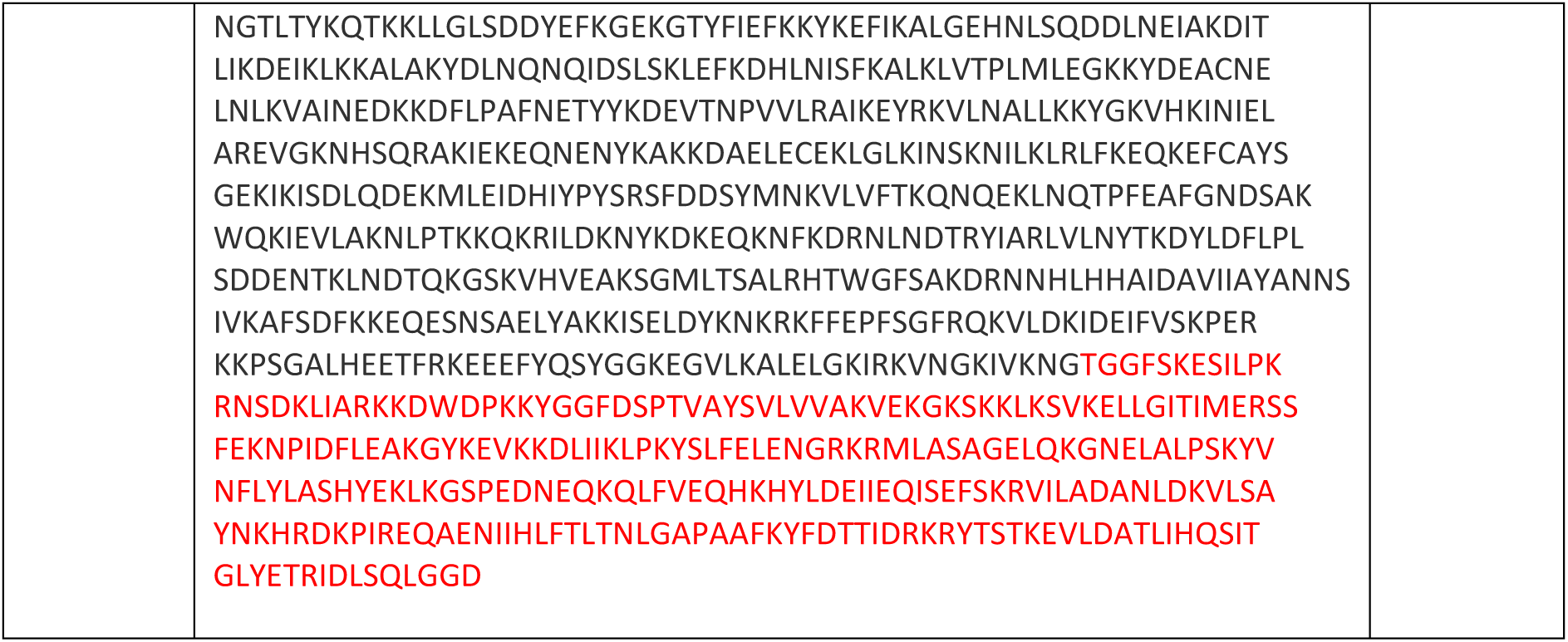
Amino acid sequence of Cas9 orthologues and of the Cas9 hybrids.

**Supplementary Table 2:**
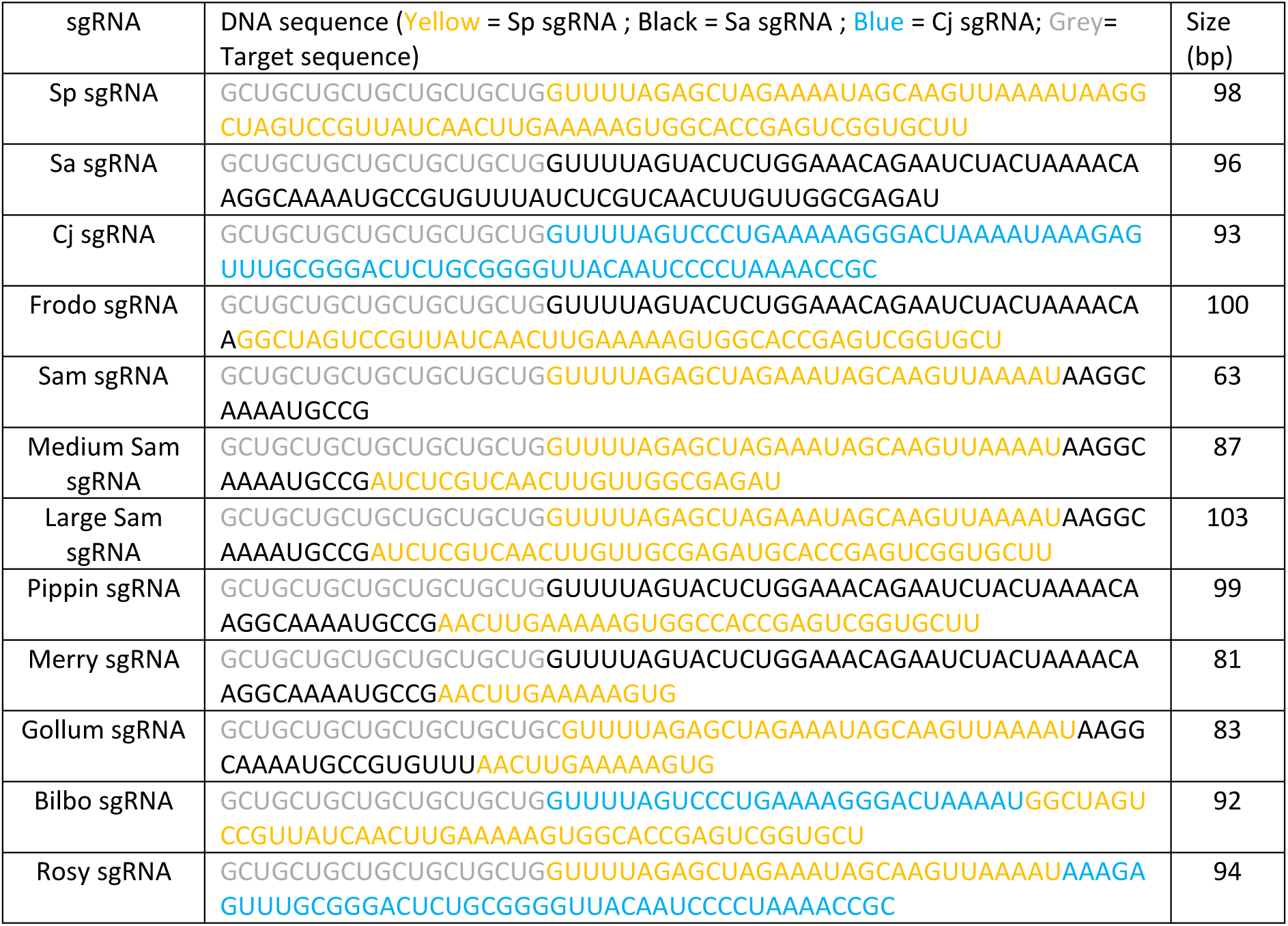
Sequence of the WT sgRNAs and hybrid sgRNAs.

**Supplementary Table 3:**
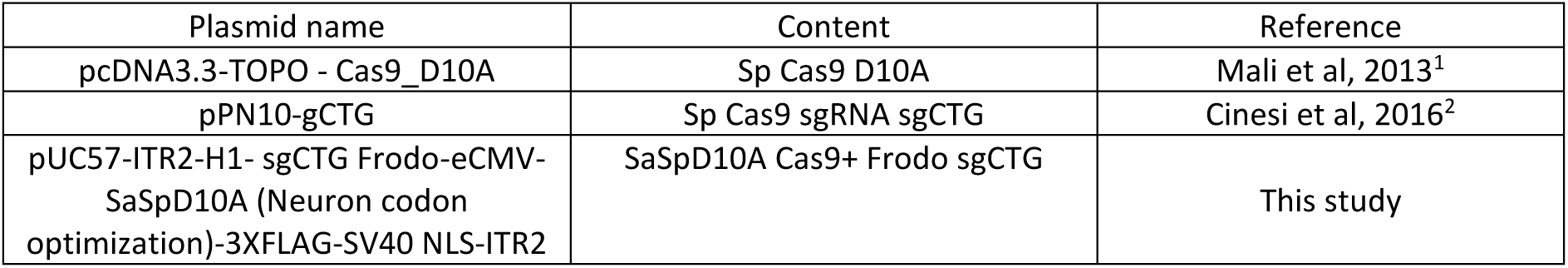
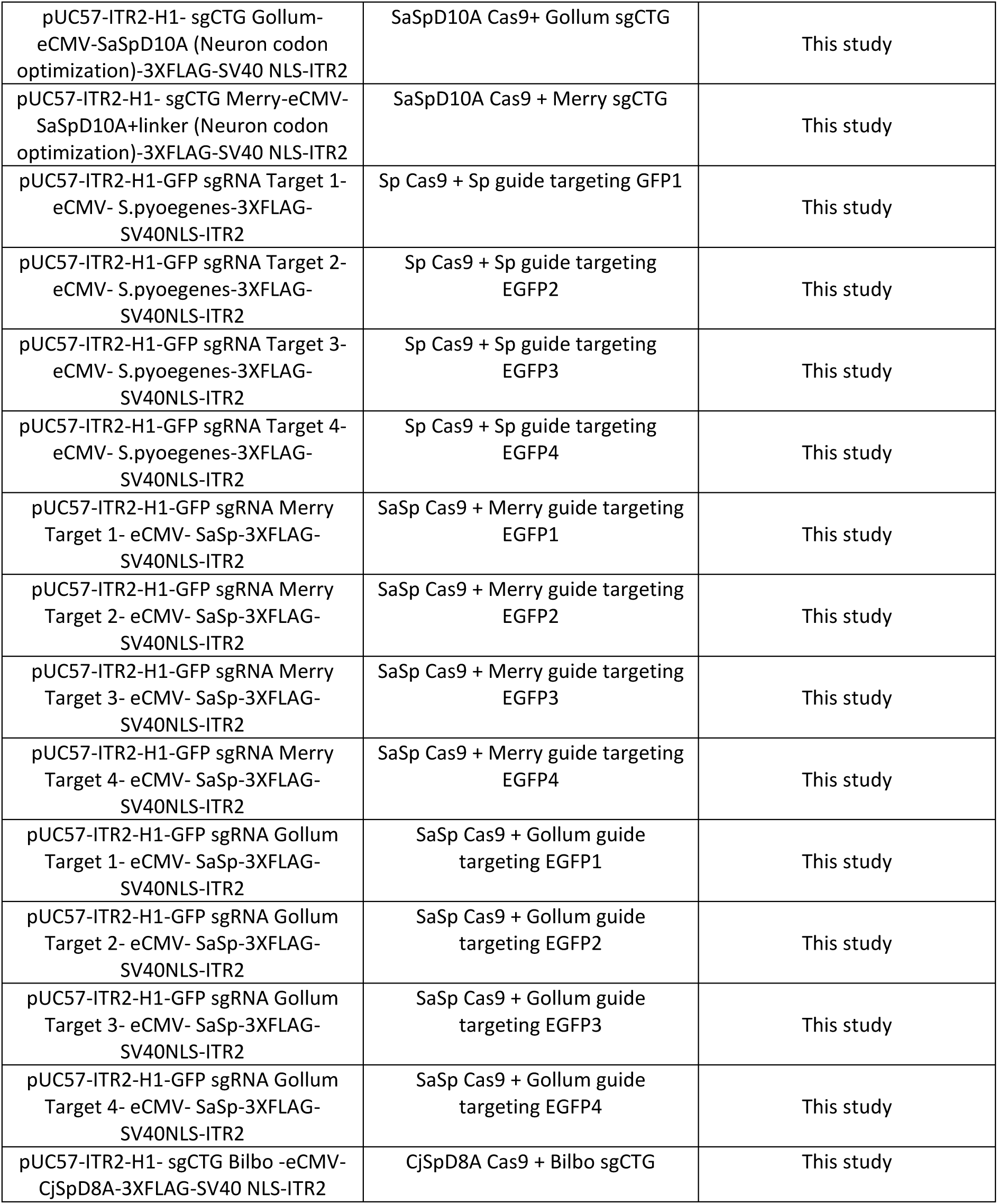
List of plasmids used.

**Supplementary Table 4:**
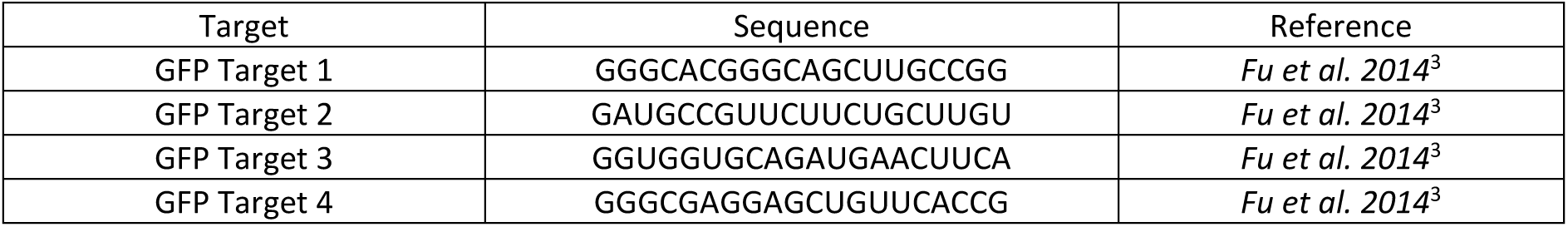
The sgRNA sequence used to target GFP.

**Supplementary Table 5:**
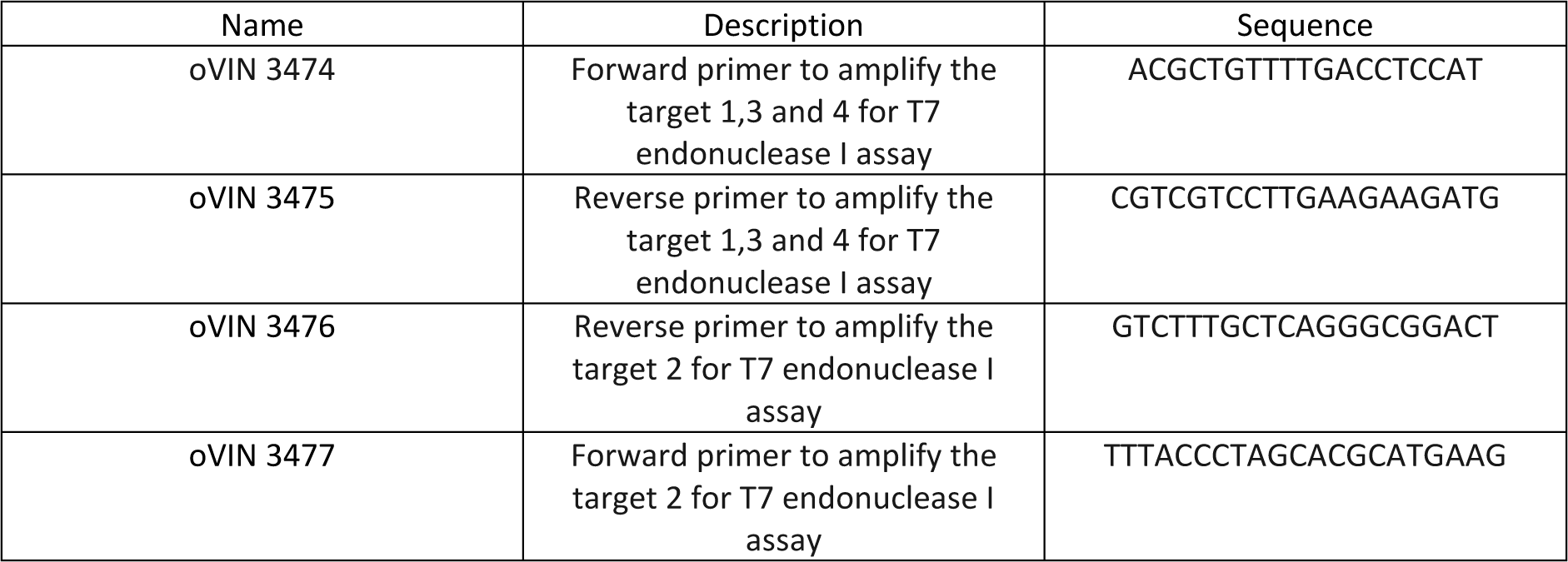
The primers used to amplify the region targeted by the GFP sgRNA.

